# Protein secretion routes in fungi are mostly determined by the length of the hydrophobic helix in the signal peptide

**DOI:** 10.1101/2025.07.30.667231

**Authors:** Tristan Sones-Dykes, Edward Wallace

## Abstract

Secreted proteins are translocated across membranes through multiple routes. In eukaryotes, secreted proteins with N-terminal signal sequences can use either the signal recognition particle (SRP) or the alternative Sec63 translocon to cross the endoplasmic reticulum membrane. Large-scale experiments on the substrates of these pathways are primarily from the model yeast *Saccharomyces cerevisiae*, but less is known about conservation of translocation pathways. Here we take a computational approach to analyse secretion signals across the fungi. Computational predictions by the Phobius model robustly separate known SRP-dependent from Sec63-dependent proteins in *S. cerevisiae*. Prior work suggested that this separation is driven by the compound hydropathy of the signal peptide’s hydrophobic helix, i.e., its length multiplied by maximum hydropathy. Instead, we find that the length of the hydrophobic helix is the major discriminator in native proteins: 8-13 amino acids for Sec63-dependent proteins and 16-27 amino acids for SRP-dependent proteins. Secreted proteins in diverse fungal species also separate into distinct populations by Phobius predictions and by the length of the hydrophobic helix. Our analysis across fungi shows that distinct functional groups of proteins, including fungal cell wall proteins and extracellular proteins, have cleaved signal peptides with short hydrophobic helices, similarly to Sec63-dependent proteins in *S. cerevisiae*. Our results suggest that the Sec63 translocon is critical for cell wall biogenesis and protein secretion in fungi, including secretion of major virulence factors in fungal pathogens of plants and animals.

**Author Summary:** Fungal cells have their “stomachs on the outside” – they secrete proteins to the environment that act as enzymes that digest macromolecules and importers of nutrients back across the cell membrane. For fungi that cause infections, the environment is the infected host animal, plant, or other fungus. Fungal cells are also protected by a cell wall, built by proteins that are secreted across the cell membrane and other proteins that use the cell’s secretory system and are then retained in the membrane. These secreted proteins are translocated across membranes by cellular machinery called translocons, and different secreted proteins use different translocons. Detailed studies of translocons in brewers’ yeast (*Saccharomyces cerevisiae*) identified the features in secreted proteins that make the protein use one translocon compared to another. Specifically, there is a helical structure near the start of secreted proteins, and current understanding is that the hydrophobicity of this helix directs the protein through the matched transposon. Here, we show that the length of the hydrophobic helix, not necessarily its maximum hydrophobicity, matters most in fungal cells: shorter helices using a translocon including the Sec63 component and longer helices using a translocon involving the “Signal Recognition Particle”. Because these translocons differ between yeast and animals, and even differ across the wide diversity of fungi, we next asked if the hydrophobic helices that direct secretion have similar properties in diverse fungi including pathogens of humans, frogs, and plants. Indeed, our computational predictions separate these these helices into short and long helix groups across fungi. The conservation of short helices in cleaved signal peptides across fungi is consistent with their using the Sec63 translocon and not the Signal Recognition Particle, which merits further investigation. Secreted proteins with cleaved signal peptides, that likely use the Sec63 translocon, include cell wall proteins, digestive enzymes, and “effectors” that manipulate the infected host to promote fungal infections.

## Introduction

Protein secretion across cell membranes is critical for cellular life. In fungi, secreted proteins include cell wall proteins that provide a scaffold to construct the cell surface; extracellular secreted proteins such as digestive enzymes that allow fungi to feed from their environment; transmembrane channels and enzymes; and secreted virulence factors and effectors in fungal pathogens of animals, plants, and other fungi [1–3]. Fungal protein secretion is also exploited when using fungi as protein production platforms [4]. Understanding the mechanisms of fungal secretion is therefore critical to understanding the fungal lifestyle, fungal pathogenesis, and biotechnological applications. More generally, secreted proteins are crucial for all eukaryotic pathogens [5]. However, detailed functional investigations of secretion mechanisms in eukaryotes have focused on a handful of model yeasts and mammalian cells, where work over the last decade has revealed unexpected complexity [6]. There is now an unmet need to better understand secretion mechanisms in non-model eukaryotes.

Secreted proteins integrate into or translocate across lipid membranes to reach their final destination, during or after being translated by cytoplasmically located ribosomes [7]. In eukaryotes, secreted proteins are first translocated across the endoplasmic reticulum (ER) membrane into the ER lumen. Here, we use “secreted” to refer to proteins that are translocated across the ER membrane, whether their final destination is intracellular, the cell surface, or extracellular, and whether or not the mature protein is retained in a membrane. The signal hypothesis states that protein secretion is specified by the protein sequence itself, in the form of an N-terminal sequence or signal peptide that directs the protein to a specific destination [8]. An ER-targeted signal peptide can be cleaved after the protein is translocated, or retained as a “signal-anchor” that first directs the protein to the ER membrane and then anchors the protein as a transmembrane helix [9]. Both kinds of signal peptides include hydrophobic helices, either as an “h-region” in a cleaved signal peptide or a transmembrane helix in a signal-anchor [9].

Secreted proteins are translocated across the ER membrane through multiple routes [10]. The signal recognition particle (SRP) was discovered in the 1980s as an RNA-protein complex, conserved in all domains of life, that binds a signal peptide as it emerges from a ribosome and interacts with a “translocon” - a transmembrane channel that permits the unfolded protein to cross the membrane co-translationally [11]. The SRP and its receptor translocate proteins with highly hydrophobic peptides and transmembrane domains [12–15]. However, the SRP does not recognise all signal peptides: in *Saccharomyces cerevisiae*, moderately hydrophobic signal peptides use an alternative Sec63-containing translocon [16]. Sec63-dependent - i.e. SRP-independent - proteins in *S. cerevisiae* constitute the majority of secreted proteins [12]. The SRP and Sec63 translocons share a core channel, called Sec61/SecY, that interacts with different proteins to recognise alternative substrates [6,17].

Signal sequences and translocons are not all functionally conserved across eukaryotes: short *S. cerevisiae* signal peptides do not function in mammalian cells [18] and the fungal Sec63 translocon contains components Sec71 and Sec72 that are apparently fungal-specific [19]. Even within fungi there is diversity in translocons, for example fungal homologs of Sec61 do not all complement the loss of *S. cerevisiae* Sec61 [20,21] while the SRP is not essential in *S. cerevisiae* but is essential in some other fungi [22,23]. Given this diversity, it is important to ask whether results on translocation mechanisms of secreted proteins can be generalised beyond model yeasts [24].

Here, we asked how the usage of translocation routes varies across fungi by undertaking a computational analysis of signal peptide sequences. First we evaluated *in silico* predictors to classify translocation routes, using *S. cerevisiae* experimental data as ground truth. We found two outputs of the Phobius algorithm [25] to be equally good at classifying protein sequences into SRP-dependent or Sec63-dependent signals: shorter 8-13 amino acid hydrophobic regions are mostly Sec63-dependent and are cleaved, while longer 16-27 amino acid regions are SRP-dependent and are retained. We then applied these effective classifiers of secreted proteins to diverse representative fungi, finding that the overall distribution of hydrophobic helix lengths is conserved.

## Results & Methods

We first evaluated signal peptide translocation route predictions in *S. cerevisiae* yeast by re-analysing published data. We directly followed the approach of Ast et al. [12], who curated a list of 1,145 *S. cerevisiae* proteins that are predicted to be translocated into the ER and then analyzed the N-terminal 60 amino acids (aa). We likewise applied Phobius [25] to the initial 60 amino acids of all annotated proteins, to generate a list of 925 proteins with predicted secretion signals within the N-terminal 60aa. Thus, every protein on this shorter list has a predicted hydrophobic helix near the N terminus, either as the h-region of a cleaved signal peptide or as a retained transmembrane helix. We compared both the length of the predicted hydrophobic helix and the maximum hydropathy of 9-aa windows on the Kyte-Doolittle scale [26].

Following Ast et al. [12], we further used ground truth experimental data to split secreted proteins into 3 groups: “Sec63-dependent” (called SRP-independent by Ast et al.) based on mislocalisation in sec72Δ cells, “SRP-dependent” based on diverse experimental datasets, and “unverified” where no clear experimental prediction is available.

Predicted hydrophobic helices show a bimodal distribution of helix lengths (Figure 1). Validated Sec63-dependent proteins have predicted 8-13 amino acid helices while SRP-dependent proteins have predicted 16-27 amino acid helices. Surprisingly, there are very few h-regions of predicted length exactly 10. Unverified proteins also have a bimodal distribution. Of 82 verified Sec63-dependent proteins, 80 have helix length 13 or less while 96 of the 107 SRP-dependent proteins have helix length of 14 amino acids or more (dashed vertical line in Figure 1). Applying a chi-squared test to this contingency table, we found that that length of the predicted helix alone can be used to classify proteins into Sec63-dependent or SRP-dependent (p < 10^−30^). Similarly, we constructed a contingency table of Phobius predictions for cleaved signal peptides or signal-anchor transmembrane helices, compared to verified Sec63-dependent or SRP-dependent categories; a chi-squared test on this contingency table also finds the prediction to be excellent (p < 10^−30^). This bimodality is not forced by the Phobius algorithm, which allows h-regions of length 6-20 amino acids within a cleaved signal peptide of indefinite total length, and transmembrane helices of length 15 to 34 amino acids, although of course the signals of shorter h-regions are included in Phobius predictions alongside the other signal peptide regions and the cleavage site [25]. The Phobius predictions, that cleaved signal peptides in *S. cerevisiae* have h-regions of length 8-13 amino acids, agrees with a previous analysis [27] that used a different algorithm, SignalP 5.0 [28].

**Figure 1.**
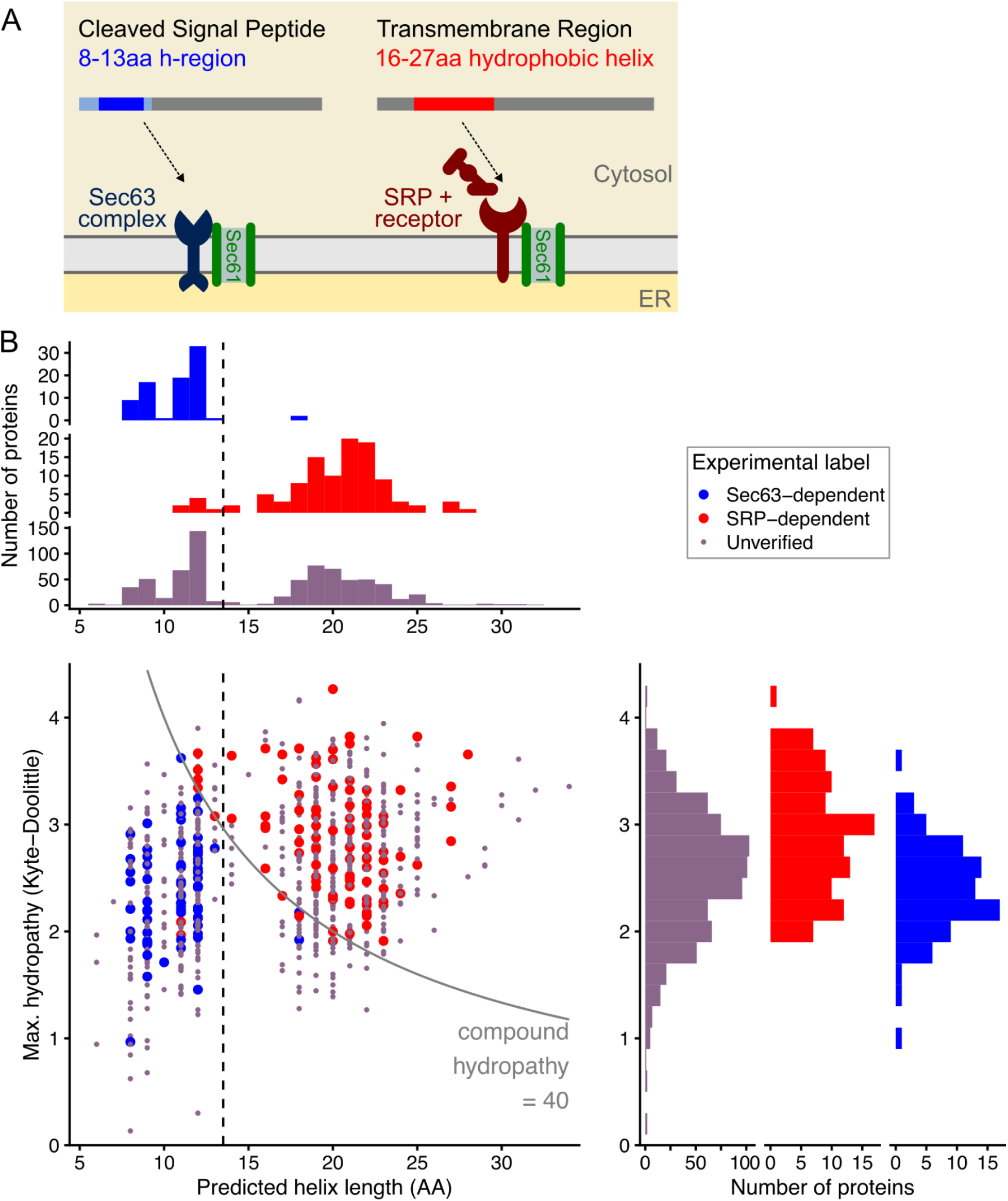
Hydrophobic helices in signal regions have a bimodal length distribution in *S. cerevisiae*, but maximum hydropathy is unimodal. A, Model of secreted proteins where cleaved signal peptides that have short hydrophobic helices (h-regions) are generally Sec63-dependent, while those with N-terminal transmembrane helices are generally SRP-dependent. B, The main panel is a scatter plot of predicted secreted proteins, showing predictions of helix length from Phobius compared to the maximum hydropathy in a 9-aa window on the Kyte-Doolittle scale. The black dashed line separates Sec63-dependent from SRP-dependent proteins by length, the grey line separates by compound hydropathy score of 40 as in Ast et al. [12]. Marginal distributions are shown: the top plot is a histogram showing the numbers of proteins by helix length, and the right plot by maximum hydropathy.

By contrast, maximum hydropathy of N-termini has a unimodal distribution (**Fig. 1**). As previously reported, maximum hydropathy for Sec63-dependent proteins is on average lower than that of SRP-dependent proteins [12], yet there is considerable overlap in their distributions. This means that maximum hydropathy is not by itself a good predictor of translocation route. Following the long-standing idea that overall hydropathy determines translocation route [16], Ast et al. computed compound hydropathy scores as the product of hydrophobic helix length and maximum hydropathy in a 9-aa window, and showed that these are bimodally distributed [12]. Repeating their analysis shows that of 82 Sec63-dependent proteins all have compound hydropathy under 40, and of 107 SRP-dependent proteins 94 have compound hydropathy over 40 (solid curve in Figure 1). A chi-squared test on this contingency table shows that compound hydropathy is an effective classifier (p < 10^−30^). However, hydrophobic helix length is a simpler metric and an equally effective classifier.

### Amino acid frequencies in hydrophobic helices differ by secretion route

We next asked if metrics beyond Kyte-Doolittle hydropathy and hydrophobic helix length, such as the Rose hydrophobicity score [29], could help to distinguish between Sec63-dependent and SRP-dependent secretion signals. We initially evaluated other hydrophobicity scales to see if they would provide a more effective classifier of translocation route. Although on average Sec63-dependent and SRP-dependent proteins have different maximum Rose hydrophobicity [29] (**Fig. S1**), we found no meaningful improvements in classification compared to using the Kyte-Doolittle scale or the predicted length of the h-region helix.

Next, we asked how the amino acid composition of hydrophobic helices differs by verified or predicted secretion route. We compared N-terminal hydrophobic helices of *S. cerevisiae* secreted proteins, using the dataset of 925 proteins analysed in Figure 1. We computed the amino acid composition of h-regions for predicted cleaved signal peptides, or of entire predicted transmembrane regions of this set of proteins. We compared 4 groups of proteins: verified Sec63-dependent proteins or SRP-dependent proteins, or those with Phobius-predicted cleaved signal peptides or retained transmembrane regions (**Fig. 2**). Verified SRP-dependent proteins and predicted N-terminal transmembrane regions have similar amino acid composition, particularly enriched in leucine. Sec63-dependent h-regions have different frequencies, with relative enrichment of serine and alanine and relative depletion of tyrosine. As the average lengths of these two groups differs, Sec63-dependent proteins have lower amino acid counts overall.

**Figure 2.**
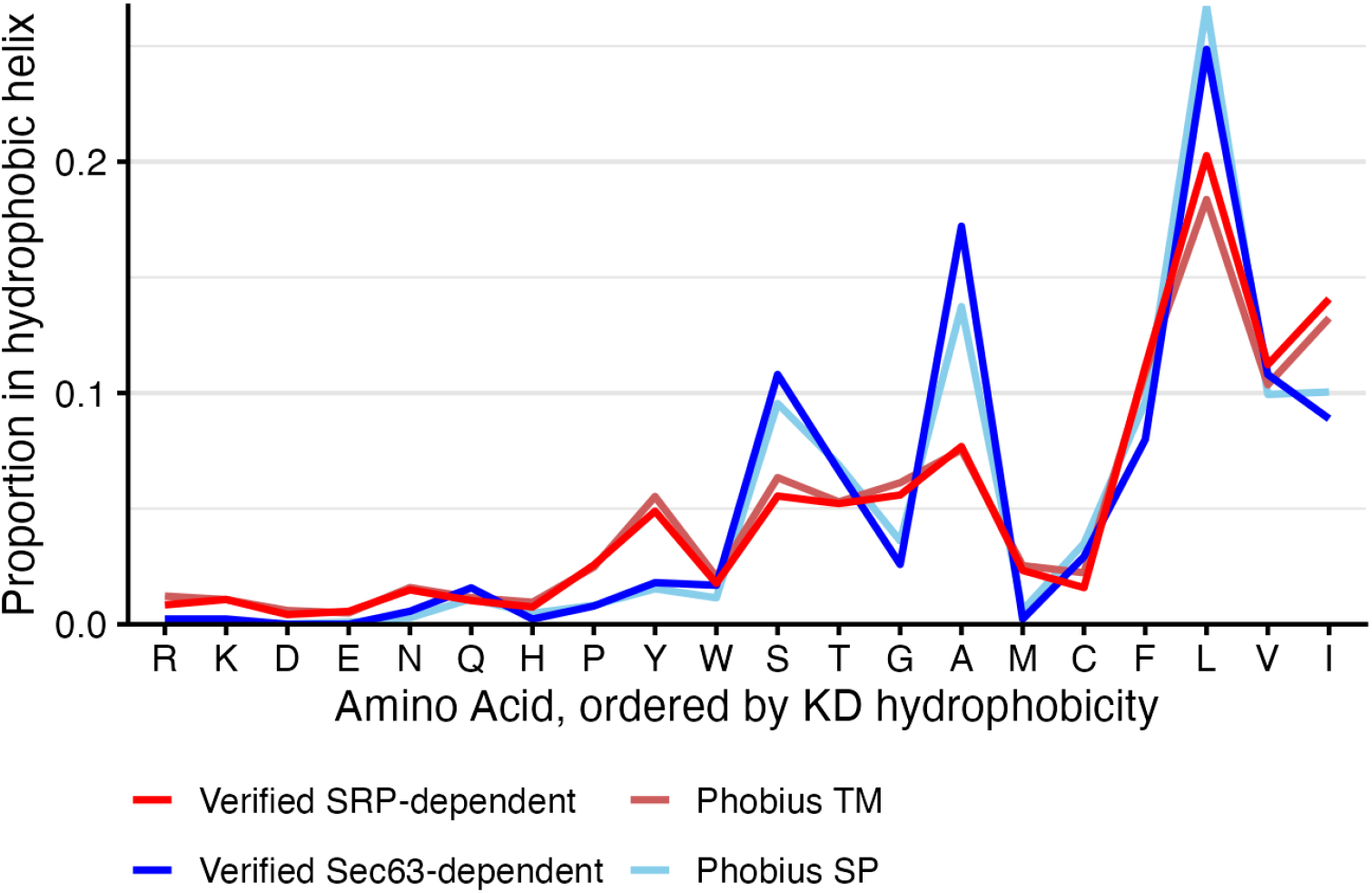
Amino acid frequencies in hydrophobic helices differ between Sec63-dependent and SRP-dependent proteins. Plot shows the proportion/frequency of amino acid usage in N-terminal hydrophobic helices of *S. cerevisiae* secreted proteins, either h-regions of cleaved signal peptides (SP) or N-terminal transmembrane helices (TM), as predicted by Phobius. Amino acids are ordered by increasing Kyte-Doolittle (KD) hydrophobicity.

Overall, this analysis of amino acid composition supports the conclusion that h-regions of verified Sec63-dependent proteins are typical of the wider set of predicted cleaved signal peptides, and likewise that transmembrane regions of verified SRP-dependent proteins are typical of the wider set of predicted N-terminal transmembrane regions. Our attempts at more detailed analysis of position-dependent amino acid frequencies yielded no clear insight. Further insight may be limited by the low absolute numbers of residues per signal peptide, for example an average of 3 leucine residues across a h-region of length 12, or 1 tyrosine in a transmembrane helix of length 20. These amino acid patterns are likely reflected in the weights embedded in the Phobius algorithm, leading to a risk of circular reasoning, and future computational work will have to address the sequence and structural determinants of translocation route in more detail.

### Comparing translocation route predictions between algorithms

To provide an independent check on our Phobius-based predictions of secretion route, we also evaluated the prediction software DeepTMHMM [30]. DeepTMHMM is a recent deep neural-network based predictor of transmembrane topology that, like Phobius, separately detects cleaved signal peptides and transmembrane helices. In the verified set of *S. cerevisiae* secretome from Ast et al. used above, DeepTMHMM cleaved signal peptide vs transmembrane helix categories also predict Sec63-dependent or SRP-dependent secretion: a chi-squared test on the contingency table of DeepTMHMM predictions compared to verified translocation route finds the prediction to be excellent (p < 10^−27^).

DeepTMHMM and Phobius mostly predict the same *S. cerevisiae* proteins as having cleaved signal peptides or retained transmembrane helices (**Fig. 3**). When both tools give a prediction, 580 out of 665 of those predictions agree; a chi-squared test on this contingency table passes (p < 10^−35^). However, some proteins are predicted in one tool but not the other, and Phobius predicts a larger number of the verified proteins as secreted.

**Figure 3.**
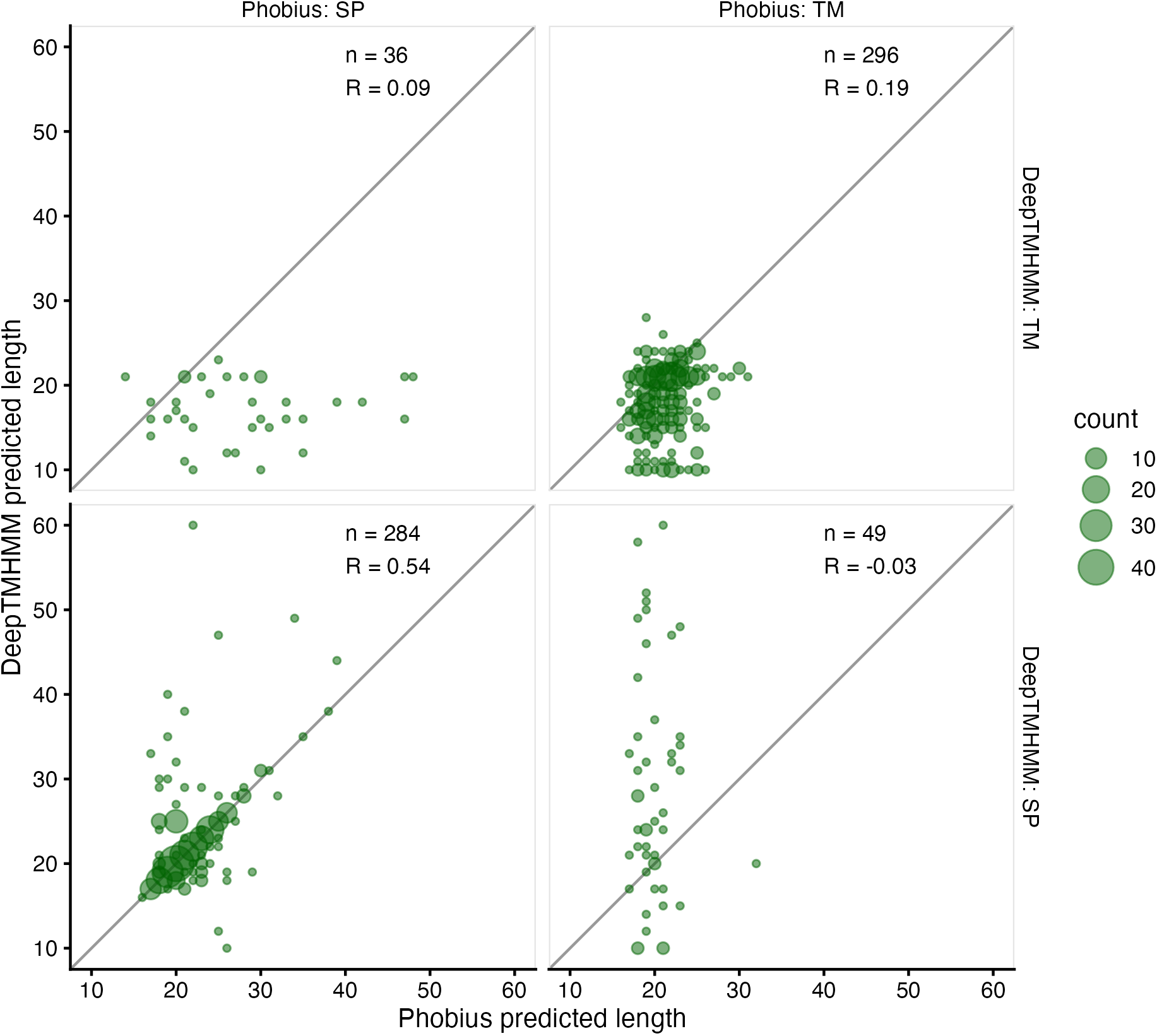
Two computational tools, DeepTMHMM and Phobius, mostly agree on their predictions of secretion signal categories and lengths. Each panel compares the predicted lengths of the entire signal region from *S. cerevisiae* proteins, and the area of each bubble is proportion to the count of proteins with that combination of predicted lengths. Panels are separated by being predicted as either cleaved signal peptide (SP) or retained signal-anchor transmembrane region (TM). This figure shows the length of the entire signal peptide, which is longer than the hydrophobic h-region, because DeepTMHMM outputs only the overall length of the signal peptide.

Unlike Phobius, DeepTMHMM does not output separate predictions of the h-region length within the cleaved signal peptide, but does predict topology for full-length proteins. Thus, we next compared predictions signal peptide class and the full signal peptide length or the N-terminal transmembrane helix length for the two tools, restricted to *S. cerevisiae* proteins for which either tool predicted such features in the initial 60 amino acids. Where their predictions agree on category, DeepTMHMM and Phobius somewhat agree on the predicted overall lengths of cleaved signal peptide (R = 0.54) (**Fig. 3**). However, correlations between predicted lengths of predicted transmembrane helices are weak (R = 0.19), and indeed some DeepTMHMM helix length predictions are too short to span a membrane. Unsurprisingly, where Phobius and DeepTMHMM predictions disagree on the category, predictions of hydrophobic helix length are uncorrelated.

We conclude that Phobius or DeepTMHMM predictions of either cleaved signal peptide vs transmembrane helix, are effective predictors of translocation route by the Sec63 complex or the SRP. Phobius and DeepTMHMM predictions generally agree on category, but not on the length of the hydrophobic helix.

### Hydrophobic helix lengths are bimodal across fungal species

If other fungi have similarly distributed signal peptide features to *S. cerevisiae*, then Phobius predictions could also predict translocation route in those fungi despite the absence of direct experimental measurements. So, we predicted signal peptides and lengths of their hydrophobic helices in a diverse selection of other fungi spanning approximately 1 billion years of fungal evolution [31]. These fungi include model organisms and common pathogens: ascomycetes (*S. cerevisiae, Candida albicans, Neurospora crassa, Magnaporthe grisea, Zymoseptoria tritici, Aspergillus fumigatus, Schizosaccharomyces pombe*), basidiomycetes (*Puccinia graminis, Ustilago maydis, Cryptococcus neoformans*), and earlier-diverging mucorales (*Rhizopus delemar)* and chytrids (*Batrachochytrium dendrobatidis*). We downloaded protein sequences from FungiDB [32] and analysed signal peptides again with Phobius [25]. We also extended the analysis to *Homo sapiens* as an outgroup, as Phobius is trained on a range of Eukaryotes including the human proteome.

We found that the distribution of predicted helix lengths for signal peptides in other fungi are highly similar to those in *S. cerevisiae*: each has a bimodal distribution of helix lengths, with almost all predicted cleaved signal peptides having h-regions of length 8-13 and almost all predicted N-terminal transmembrane helices having length 16-27 (**Fig. 4**). The numbers of proteins in each category vary widely, for example *P. graminis* has roughly 10 times the number of proteins with cleaved signal peptides as *S. pombe*. There are some minor differences in the distribution, for example the early-diverging chytrid *B. dendrobatis* has more h-regions of lengths 11 and 13 relative to length 12 that could indicate differences in preferred h-regions. Overall, the conserved bimodal distributions are consistent with cleaved signal peptides with short hydrophobic helical h-region being categorically different from retained signal-anchors that have longer transmembrane regions. This suggests that their distinct translocation routes may also be conserved across fungi.

**Figure 4.**
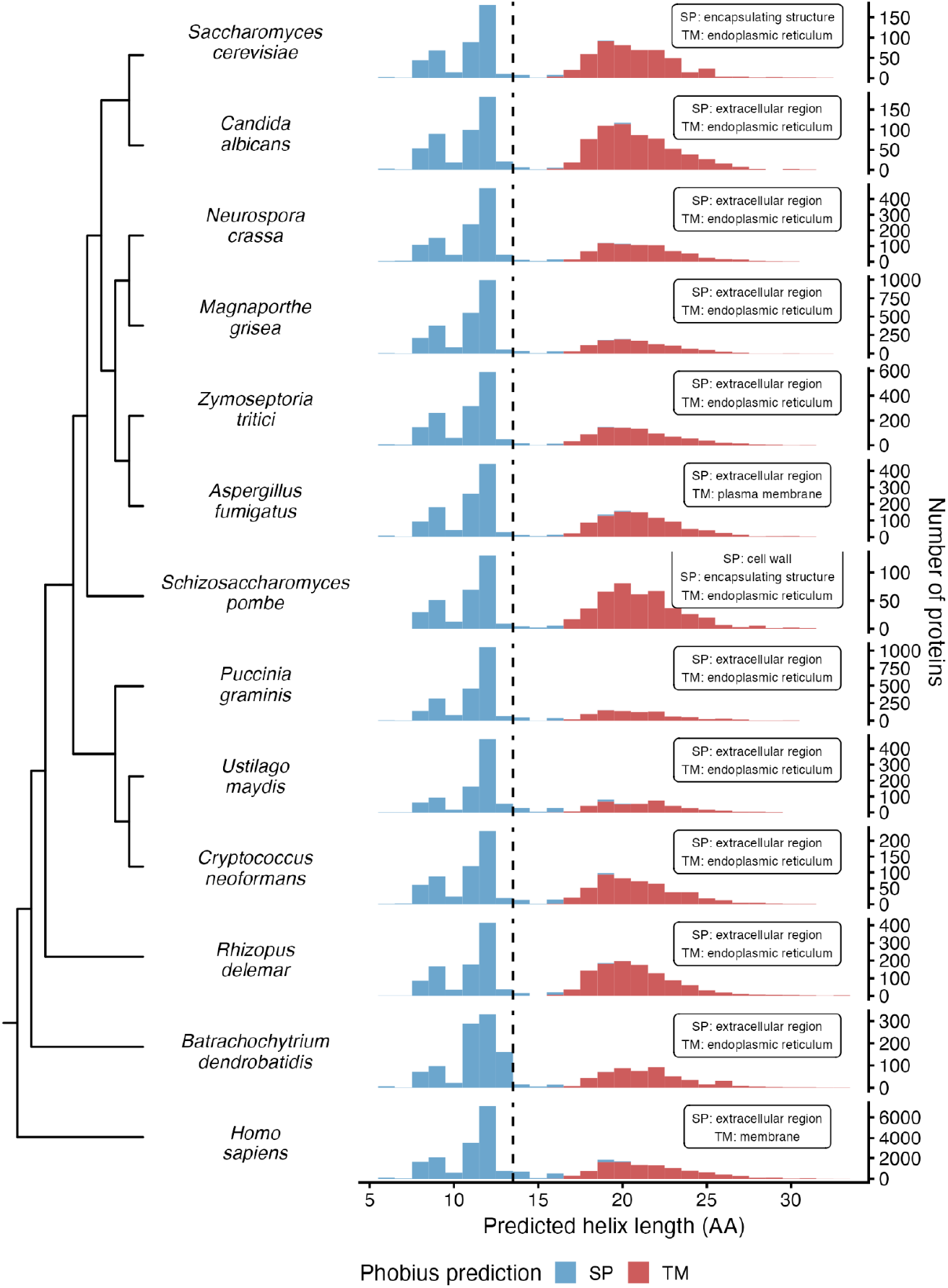
Predicted hydrophobic helices in signal peptides have a bimodal length distribution across fungi. Left panel shows cladogram of 12 included fungal species and Homo sapiens, right panels show a histogram of the number of proteins with each helix length predicted by Phobius, coloured as the h-region of a cleaved signal peptide (SP) or as the transmembrane helix of a retained signal-anchor (TM). The dashed line indicates the threshold at 13 amino acids, as in Figure 1. The y-axis scales are different in each panel, reflecting differences in the total number of secreted proteins between species. The inset text shows the GOSlim term with the lowest p-value for each predicted group, except for *S. pombe* where 2 terms had the same p-value. We abbreviated “external encapsulating structure” to “encapsulating structure”.

To understand the functional properties of proteins with predicted cleaved signal peptides or signal-anchor transmembrane regions, we performed a gene ontology (GO) analysis using FungiDB [32]. As expected, across fungi, proteins with predicted cleaved signal peptides are enriched in GO terms for cell wall (GO:0005618), external encapsulating structure (GO:0030312), and extracellular region (GO:0005576). Both cleaved signal peptides and predicted transmembrane regions are enriched for GO terms for endoplasmic reticulum (GO:0005783), plasma membrane (GO:0005886), and in some cases vacuole (GO:0005773) and Golgi apparatus (GO:0005794). In all cases the p-values for GO enrichment are below 10^−10^. This analysis supports conservation of protein functions within each category, despite the limits of GO analysis in non-model fungi.

The analysis of the *Homo sapiens* proteome supports our findings in fungi, despite the larger proteome size and higher number of Phobius-predicted proteins (**Fig. 4)**. Predicted h-region lengths in humans had a similar distribution. Also GO terms from PANTHER GO-slim [33] match with those found by FungiDB for fungi species. For mammalian proteins, 4 leucines in a N-terminal transmembrane region enable it to stably insert into the endoplasmic reticulum membrane [15]. This agrees with our amino-acid frequency analysis of the entire Phobius-predicted fungal protein set (**Fig. 2**).

Fitting a logistic regression model to classify verified *S. cerevisiae* proteins, based on a binary predictor of at least 4 leucines in the Phobius predicted window, finds it to be highly significant (p < 10^−8^). Calculating the odds-ratio shows that the predicted secondary structures are approx 8.9 times more likely to be transmembrane regions, given at least 4 leucines.

This consolidates the relationship between predicting secretion pathways in fungi and mammals. A classification method based on length, that is verified using fungal proteomes, produces the same patterns in *Homo sapiens*; and a result from mammal protein experiments can be used to create a classification heuristic in fungal proteomes.

## Discussion

### Hydrophobic helix length determines translocation route of signal peptides in Fungi

This work extends and simplifies previous results on how N-terminal sequences specify the route of translocation into the ER. We used curated experimental data in *S. cerevisiae* and validated computational prediction algorithms to show that, overwhelmingly, Sec63-dependent proteins have cleaved signal peptides while SRP-dependent proteins have retained transmembrane signal-anchor peptides. Consistent with previous observations in fungi and mammals, cleaved signal peptides have shorter hydrophobic helices than signal anchors [25], and fewer leucines [15]. By contrast, the maximum hydrophobicity of the predicted signal region is not a strong discriminator of the different translocation pathways.

Our analysis here uses primarily the Phobius algorithm, which predicts the overwhelming majority of fungal cleaved signal peptides to have h-regions of length 13 or less despite the model allowing cleaved signal peptides to have length up to 20 amino acids [25]. This analysis agrees with predictions of h-region lengths from SignalP 5.0 [28] and of overall signal peptide length from DeepTMHMM (**Fig. 3**). Further structural data would be needed to test the differing algorithms’ predictions of h-region length, including Phobius’ prediction of a lack of h-regions of length 10, for which we cannot think of a good explanation.

Our analysis agrees with diverse observations arguing that the fungal Sec63 complex efficiently translocates signal peptides with short hydrophobic helices, and that this is not exactly conserved in mammals. Mutational studies showed that shortening the signal sequence of *S. cerevisiae* invertase maintained its secretion independent of SRP function [34]. Furthermore, shortened invertase signal peptides were not secreted in mammalian cells and were not recognised by SRP [34]. The Sec63-dependent signal peptide of yeast carboxypeptidase Y does not support translocation in mammalian cells [18]. In contrast, shortening h-regions of Sec63-dependent signal peptides can lead to more efficient secretion [27]. The apparent preference of the fungal Sec63 translocon for shorter h-regions may relate to the presence of fungal-specific components on its cytoplasmic surface, homologs of Sec71 and Sec72 in *S. cerevisiae* [17]. However, Phobius’ predictions of equally short h-regions in human proteins (**Fig. 4**) suggests that length alone may not be a complete explanation, and one report argues that h-regions of Sec62/Sec63 dependent signal peptides in humans are longer, with a mode of 15 amino acids [35].

The critical role played by the length of hydrophobic helices in targeting is supported by structural evidence. Signal-anchors are retained transmembrane helices and have an average length of 24 amino acids [36], which is appropriate to span a lipid bilayer of width 35-40Å. Structures of the translocating SRP-Sec61 complex show helices of length 17 or 19 amino acids interacting with Sec61 (PDB: 4CG6 and 3JC2) [37,38]. By contrast, cleaved signal peptides do not need to span a membrane. A structure of the active Sec63-Sec61 translocon includes a hydrophobic helix in the signal peptide of length ∼8 amino acids interacting with Sec61 (PDB: 7AFT) [39].

Experimental evidence shows that some signal peptides can be recognised by multiple translocons [12]. Mutational studies show that increasing the hydrophobicity of Sec63-dependent signal peptides is sufficient for them to be recognised by SRP [16]. Indeed, some *S. cerevisiae* proteins with short but highly hydrophobic h-regions are reported as SRP dependent [12]. Thus, although signal regions in native fungal secreted proteins have a bimodal distribution of properties corresponding to two major translocation routes, sequences with intermediate properties can still be translocated by one or both of the translocons.

Overall, the evidence argues that the Sec63 translocon mostly recognizes short hydrophobic helices with a length of 8-13 amino acids in native proteins. Describing Sec63-dependent signal peptides as “moderately hydrophobic” or “marginally hydrophobic” subtly misses the point: the h-regions of cleaved signal peptides are generally short.

### The importance of the Sec63 translocon in fungi

Our analysis argues that secreted proteins in fungi and beyond are separated into distinct subsets that depend on distinct translocons. These subsets consist of structurally and functionally distinct groups that are conserved across the fungal kingdom. In other fungi, as in *S. cerevisiae*, proteins with retained signal-anchors that are long transmembrane helices are very likely to use the SRP. By contrast, secreted cell wall proteins and others with cleaved signal peptides and shorter hydrophobic helices are likely to use the Sec63 translocon.

Our results suggest that the regulation and function of the Sec63 translocon is critical for cell wall biogenesis and protein secretion across fungi. Secreted proteins with cleaved signal peptides include major secreted virulence factors in fungal pathogens of plants and animals such as *C. albicans* candidalysin and secreted aspartyl proteases [3]. Predicted cleaved signal peptides are already used to computationally identify candidate secreted effector proteins in plant fungal pathogens that might suppress host immunity or modify host cellular activities [40,41]. Indeed computational screens of conserved signal peptide-containing proteins can be an efficient way to discover virulence factors [42]. Thus, investigating fungal secretion mechanisms, including translocation routes and their regulation, will be critical to understand fungal pathogenesis.

Further work is needed to experimentally validate our computational predictions. In *S. cerevisiae*, partial loss-of-function mutants of Sec62, Sec63, and the non-essentiality of Sec71 and Sec72, facilitated genetic analysis of their substrates [16]. These results were validated by a combination of large-scale genetic screens, functional genomics, and detailed experiments with signal peptide swaps in reporter genes [12–14]. Similar experiments in diverged fungi could directly measure the substrates of alternative secretion pathways. We speculate that there may be yet more complexity in secretion pathways in other fungi, especially in early-diverging fungi and in multicellular fungi with complex lifestyles and large secretomes.

## Data availability

All data, data analysis code, and figures, are available at https://github.com/TristanSones-Dykes/TMSP_Pub. We will mint a doi for archiving when we submit for publication.

## Acknowledgments

We thank Atlanta Cook for extensive discussions and critical feedback throughout the project. We thank Maya Schuldiner for inspiration and encouragement. We thank Alex Cope and Wallace lab members, for comments and feedback on the manuscript. E.W.J.W. would also like to thank his co-authors, lab members, and Liz Ballou, for tolerating his enthusiasm about signal peptides over the course of this project. E.W.J.W was supported by Wellcome Sir Henry Dale Fellowship [208779/Z/17/Z], and T.S-D. was supported by a Wellcome BVS [220234].

---

**Figure S1.**
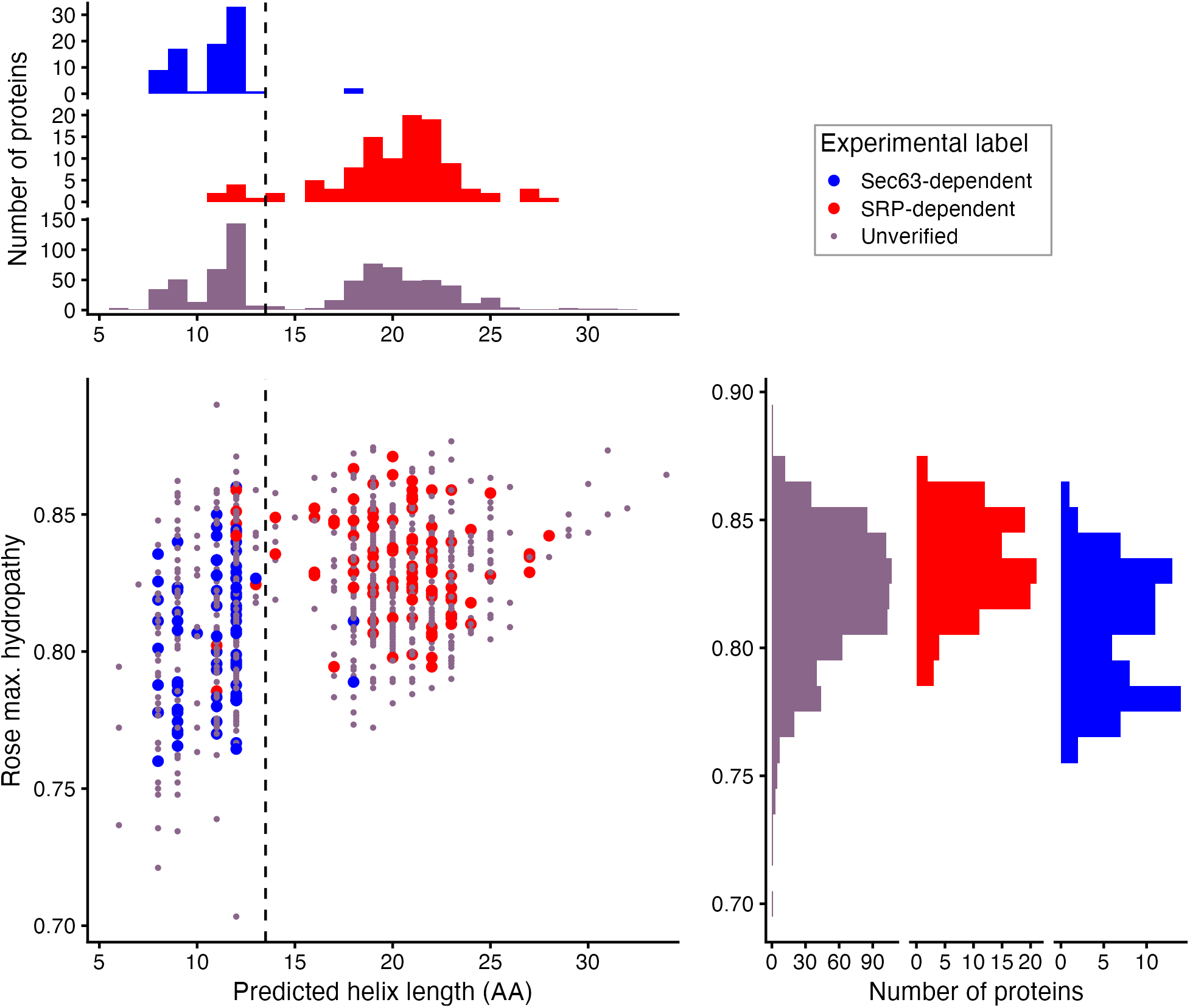
related to figure 1. Hydrophobic helices in signal regions in *S. cerevisiae*, have a wider distribution of maximum hydropathy calculated on the Rose scale. This figure is similar to Figure 1, except calculating the Rose hydrophobicity of 9 aa window (y axis) instead of the Kyte-Doolittle hydrophobicity.

## Notes

https://github.com/TristanSones-Dykes/TMSP_Pub

### Competing Interest Statement

The authors have declared no competing interest.

## References

1. Gow NAR, Lenardon MD. Architecture of the dynamic fungal cell wall. Nat Rev Microbiol. 2022. doi:10.1038/s41579-022-00796-9

2. Ellis JG, Dodds PN, Lawrence GJ. The role of secreted proteins in diseases of plants caused by rust, powdery mildew and smut fungi. Curr Opin Microbiol. 2007;10: 326–331.

3. Brunke S, Mogavero S, Kasper L, Hube B. Virulence factors in fungal pathogens of man. Curr Opin Microbiol. 2016;32: 89–95.

4. Conesa A, Punt PJ, van Luijk N, van den Hondel CA. The secretion pathway in filamentous fungi: a biotechnological view. Fungal Genet Biol. 2001;33: 155–171.

5. Haldar K, Kamoun S, Hiller NL, Bhattacharje S, van Ooij C. Common infection strategies of pathogenic eukaryotes. Nat Rev Microbiol. 2006;4: 922–931.

6. Aviram N, Schuldiner M. Targeting and translocation of proteins to the endoplasmic reticulum at a glance. J Cell Sci. 2017;130: 4079–4085.

7. Rapoport TA. Protein translocation across the eukaryotic endoplasmic reticulum and bacterial plasma membranes. Nature. 2007;450: 663–669.

8. Blobel G, Walter P, Chang CN, Goldman BM, Erickson AH, Lingappa VR. Translocation of proteins across membranes: the signal hypothesis and beyond. Symp Soc Exp Biol. 1979;33: 9–36.

9. Martoglio B, Dobberstein B. Signal sequences: more than just greasy peptides. Trends Cell Biol. 1998;8: 410–415.

10. Ast T, Schuldiner M. All roads lead to Rome (but some may be harder to travel): SRP-independent translocation into the endoplasmic reticulum. Crit Rev Biochem Mol Biol. 2013;48: 273–288.

11. Pool MR. Signal recognition particles in chloroplasts, bacteria, yeast and mammals (review). Mol Membr Biol. 2005;22: 3–15.

12. Ast T, Cohen G, Schuldiner M. A network of cytosolic factors targets SRP-independent proteins to the endoplasmic reticulum. Cell. 2013;152: 1134–1145.

13. Costa EA, Subramanian K, Nunnari J, Weissman JS. Defining the physiological role of SRP in protein-targeting efficiency and specificity. Science. 2018;359: 689–692.

14. Jan CH, Williams CC, Weissman JS. Principles of ER cotranslational translocation revealed by proximity-specific ribosome profiling. Science. 2014;346: 1257521.

15. Hessa T, Kim H, Bihlmaier K, Lundin C, Boekel J, Andersson H, et al. Recognition of transmembrane helices by the endoplasmic reticulum translocon. Nature. 2005;433: 377–381.

16. Ng DT, Brown JD, Walter P. Signal sequences specify the targeting route to the endoplasmic reticulum membrane. J Cell Biol. 1996;134: 269–278.

17. Rapoport TA, Li L, Park E. Structural and Mechanistic Insights into Protein Translocation. Annu Rev Cell Dev Biol. 2017;33: 369–390.

18. Bird P, Gething MJ, Sambrook J. Translocation in yeast and mammalian cells: not all signal sequences are functionally equivalent. J Cell Biol. 1987;105: 2905–2914.

19. Schlegel T, Mirus O, von Haeseler A, Schleiff E. The tetratricopeptide repeats of receptors involved in protein translocation across membranes. Mol Biol Evol. 2007;24: 2763–2774.

20. Broughton J, Swennen D, Wilkinson B, Joyet P, Gaillardin C, Stirling C. Cloning of SEC61 homologues from Schizosaccharomyces pombe and Yarrowia lipolytica reveals the extent of functional conservation within this core component of the ER translocation machinery. J Cell Sci. 1997;110 (Pt 21): 2715–2727.

21. de la Rosa JM, Ruiz T, Fonzi WA, Rodríguez L. Analysis of heterologous expression of Candida albicans SEC61 gene reveals differences in Sec61p homologues related to species-specific functionality. Fungal Genet Biol. 2004;41: 941–953.

22. Brennwald P, Liao X, Holm K, Porter G, Wise JA. Identification of an essential Schizosaccharomyces pombe RNA homologous to the 7SL component of signal recognition particle. Mol Cell Biol. 1988;8: 1580–1590.

23. He F, Yaver D, Beckerich JM, Ogrydziak D, Gaillardin C. The yeast Yarrowia lipolytica has two, functional, signal recognition particle 7S RNA genes. Curr Genet. 1990;17: 289–292.

24. Delic M, Valli M, Graf AB, Pfeffer M, Mattanovich D, Gasser B. The secretory pathway: exploring yeast diversity. FEMS Microbiol Rev. 2013;37: 872–914.

25. Käll L, Krogh A, Sonnhammer ELL. A combined transmembrane topology and signal peptide prediction method. J Mol Biol. 2004;338: 1027–1036.

26. Kyte J, Doolittle RF. A simple method for displaying the hydropathic character of a protein. J Mol Biol. 1982;157: 105–132.

27. Xue S, Liu X, Pan Y, Xiao C, Feng Y, Zheng L, et al. Comprehensive Analysis of Signal Peptides in Saccharomyces cerevisiae Reveals Features for Efficient Secretion. Adv Sci. 2023;10: e2203433.

28. Almagro Armenteros JJ, Tsirigos KD, Sønderby CK, Petersen TN, Winther O, Brunak S, et al. SignalP 5.0 improves signal peptide predictions using deep neural networks. Nat Biotechnol. 2019;37: 420–423.

29. Rose GD, Geselowitz AR, Lesser GJ, Lee RH, Zehfus MH. Hydrophobicity of amino acid residues in globular proteins. Science. 1985;229: 834–838.

30. Hallgren J, Tsirigos KD, Pedersen MD, Armenteros JJA, Marcatili P, Nielsen H, et al. DeepTMHMM predicts alpha and beta transmembrane proteins using deep neural networks. bioRxiv. 2022. p. 2022.04.08.487609. doi:10.1101/2022.04.08.487609

31. Li Y, Steenwyk JL, Chang Y, Wang Y, James TY, Stajich JE, et al. A genome-scale phylogeny of the kingdom Fungi. Curr Biol. 2021;31: 1653–1665.e5.

32. Basenko EY, Pulman JA, Shanmugasundram A, Harb OS, Crouch K, Starns D, et al. FungiDB: An Integrated Bioinformatic Resource for Fungi and Oomycetes. J Fungi (Basel). 2018;4. doi:10.3390/jof4010039

33. Mi H, Muruganujan A, Ebert D, Huang X, Thomas PD. PANTHER version 14: more genomes, a new PANTHER GO-slim and improvements in enrichment analysis tools. Nucleic Acids Res. 2019;47: D419–D426.

34. Rothe C, Lehle L. Sorting of invertase signal peptide mutants in yeast dependent and independent on the signal-recognition particle. Eur J Biochem. 1998;252: 16–24.

35. Schorr S, Nguyen D, Haßdenteufel S, Nagaraj N, Cavalié A, Greiner M, et al. Identification of signal peptide features for substrate specificity in human Sec62/Sec63-dependent ER protein import. FEBS J. 2020;287: 4612–4640.

36. Saidijam M, Azizpour S, Patching SG. Comprehensive analysis of the numbers, lengths and amino acid compositions of transmembrane helices in prokaryotic, eukaryotic and viral integral membrane proteins of high-resolution structure. J Biomol Struct Dyn. 2018;36: 443–464.

37. Gogala M, Becker T, Beatrix B, Armache J-P, Barrio-Garcia C, Berninghausen O, et al. Structures of the Sec61 complex engaged in nascent peptide translocation or membrane insertion. Nature. 2014;506: 107–110.

38. Voorhees RM, Hegde RS. Structure of the Sec61 channel opened by a signal sequence. Science. 2016;351: 88–91.

39. Weng T-H, Steinchen W, Beatrix B, Berninghausen O, Becker T, Bange G, et al. Architecture of the active post-translational Sec translocon. EMBO J. 2021;40: e105643.

40. Seong K, Krasileva KV. Prediction of effector protein structures from fungal phytopathogens enables evolutionary analyses. Nat Microbiol. 2023;8: 174–187.

41. Derbyshire MC, Raffaele S. Surface frustration re-patterning underlies the structural landscape and evolvability of fungal orphan candidate effectors. Nat Commun. 2023;14: 5244.

42. Schuster M, Schweizer G, Reißmann S, Happel P, Aßmann D, Rössel N, et al. Novel Secreted Effectors Conserved Among Smut Fungi Contribute to the Virulence of Ustilago maydis. Mol Plant Microbe Interact. 2024;37: 250–263.

